# Reconstruction of Full-length scFv Libraries with the Extended Range Targeted Sequencing Method

**DOI:** 10.1101/2022.05.10.491248

**Authors:** Christopher Wei, Sarah Criner, Bharat Sridhar, Ryan Shultzaberger, Timothy Looney, Martin M Fabani, Eli N Glezer

## Abstract

Single chain fragment variable (scFv) phage display libraries of randomly paired VH-VL antibody domains are a powerful and widely adopted tool for the discovery of antibodies of a desired specificity. Characterization of full length VH-VL constructs using synthetic long read assemblies of short read next-generation sequencing data has emerged as a powerful approach to identify antibody candidates with greater speed and sensitivity than classical screening methods. Here we introduce a new version of the synthetic long read approach, which we denote the Extended Range Targeted Sequencing method. We apply the method to demonstrate accurate and high throughput analysis of full-length VH-VL constructs from a commercial scFv combinatorial display library.

## Introduction

Next-generation Sequencing (NGS) is paving the way towards personalized medicine, not only by enabling vast breakthroughs in understanding the onset of disease (Ng et al. 2009), disease progression (Onidani et al. 2019) and treatment (Orlov-Slavu et al. 2021), but also from its impact on other developing technologies (Shahid et al. 2021). Single chain fragment variable (scFv) phage display libraries of randomly paired VH-VL antibody domains are a powerful and widely adopted tool for the discovery of antibodies of a desired specificity. Traditional phage display workflows employ multiple rounds of antigen panning to enrich for one or several dominating antibody clones, which are then further screened by ELISA before Sanger sequencing. Limitations of this method include the inability to analyze targets that do not readily elicit dominating clones (e.g., G-protein-coupled receptors, Jo et al. 2016), inability to identify subdominant but relevant binders, and the need to conduct multiple time-consuming rounds of selection to identify antibody candidates. Due to its high throughput and relatively low cost, NGS has been touted as an ideal tool to characterize scFv libraries (Glanville et al. 2015, Rouet et al. 2018). However, due to read length limitations, this application has been predominantly served by more expensive and generally less accurate long-read sequencing technologies (Hemadou et al. 2017, Han et al. 2018, Nannini et al. 2021). Various library preparation methods have been developed to overcome the current read length limitations in NGS, and have been demonstrated for both scFv (Turchaninova et al.2016, Cole et al.2016, Burke et al. 2016) and 16S RNA sequencing (Callahan et al. 2021, Karst et al. 2018).

In this work we introduce a novel library preparation approach, the Extended Range Targeted Sequencing method (XR/T-Seq™ method), that enables long molecule reconstruction from standard paired 2×150bp reads. We demonstrate the utility of this method by characterizing the molecular diversity, composition, and general properties of a commercial scFv phage combinatorial antibody display library.

## Results

The work presented here is aimed at demonstrating a simple but powerful tool to enable the reconstruction of ~1-3Kb long fragments from short read technology. The method, shown in Figure 1A, relies on the introduction of Unique Molecular Identifiers (UMI) at invariant positions along single molecule templates. 12bp barcodes are embedded in linear and stem-loop probes that, after sequence-specific binding, are extended and ligated to neighboring probes by a combination of a non-strand displacing DNA Polymerase and a fast DNA Ligase. Following extension and ligation, UMI-containing full-length products are quantified, diluted to a desired input and amplified by PCR. This limited complexity library (i.e., diluted) is then processed and sequenced by standard NGS protocols. To carry out molecule sequence reconstruction, we developed a companion bioinformatic pipeline (Figure 1B) consisting of i) UMI identification and collapse, ii) UMI co-occurrence analysis and read grouping, iii) k-mer based read filtering, and iv) assembly.

**Figure 1.**
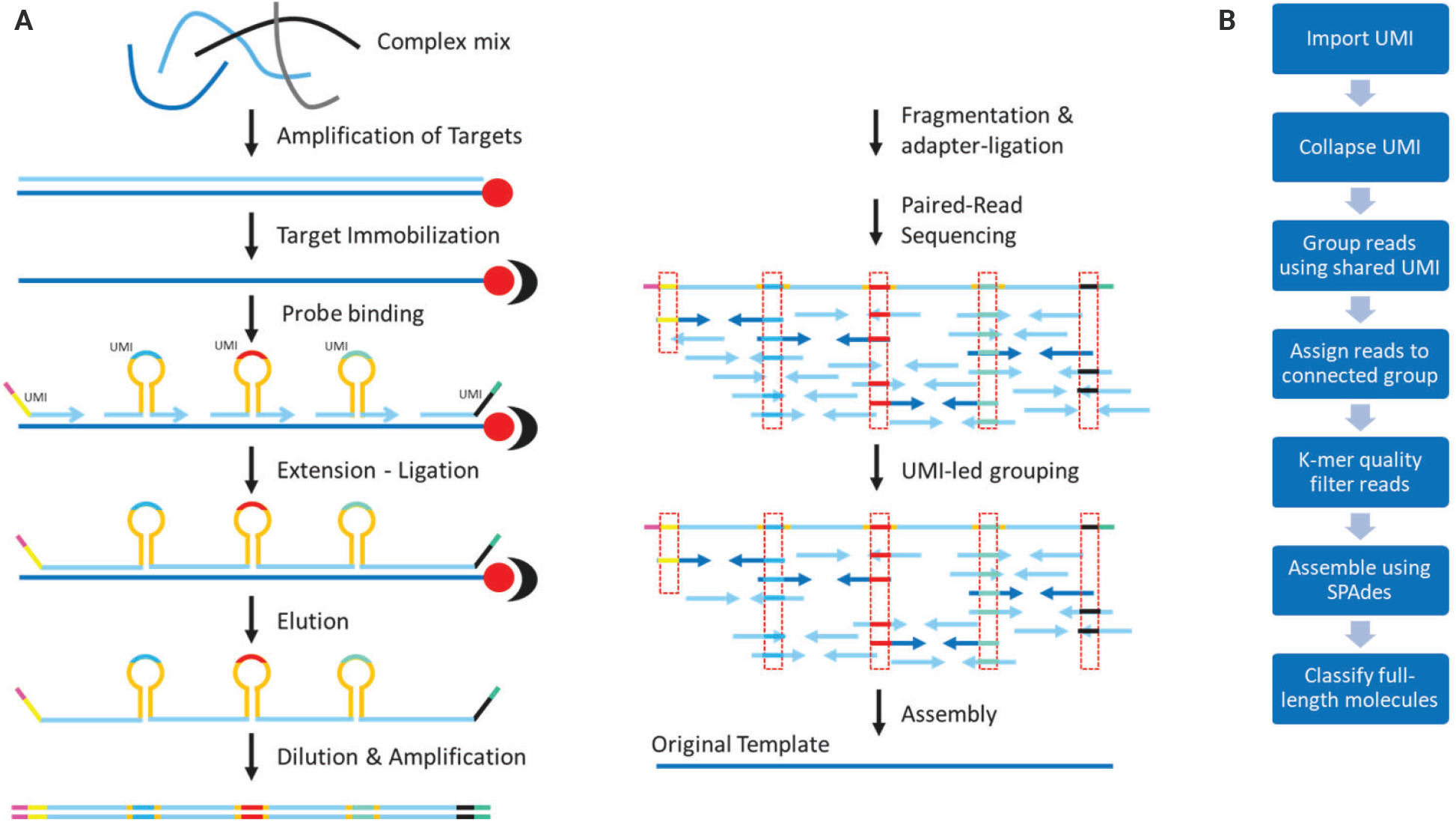
The XR/T-Seq library preparation workflow. A. Overview of the XR/T-Seq method, including the library preparation workflow for single molecule tagging and synthetic long read assembly. Single molecule barcodes are introduced into defined positions within a region of interest (typically, invariant windows within an otherwise mutable region) by hybridization and extension-ligation of UMI containing probes. The integrated strand containing UMIs and complementary template sequences are amplified, fragmented, then sequenced using paired 2×150bp reads. B. Following sequencing, UMIs are extracted and co-occurrence analysis identifies UMI groups derived from the same molecule. Finally, UMI groups are used to cluster reads, which undergo k-mer based filtering prior to assembly.

In order to demonstrate the utility of the XR/T-Seq method, we decided to apply it to the determination of molecular diversity in a commercial scFv phage display library (Figure 2) consisting of randomly paired VH and VL domains from IgG-expressing B cells derived from a pool of healthy donors. The library VH and VL domains are separated by an invariant 45bp flexible linker sequence, for a total insert size of ~0.8-1Kb. This model system is an ideal testing platform for XR/T-Seq due to its intrinsic characteristics: the diversity of the library is very high (>10^13^ unique members), the length of the templates is beyond the read length of short read sequencing technologies, and the sequence is difficult to reconstruct by simple alignment of short reads.

**Figure 2.**
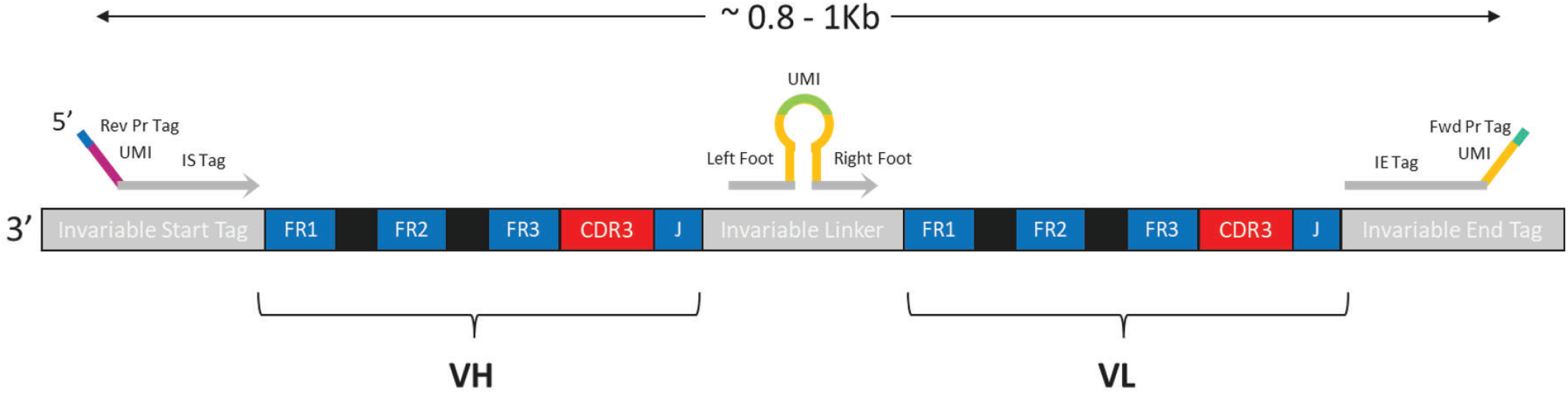
XR/T-Seq method UMI tagging strategy for scFv library analysis. The XR/T-Seq method was applied to analyze a commercial scFv phage display library consisting of randomly paired VH and VL domains from IgG-expressing B cells derived from a pool of healthy donors. UMIs are introduced into terminal ends and the invariant linker region of the ~950bp VH-VL construct via UMI tagged external primers in conjunction with a linker-targeting hairpin loop adapter. Primers and the hairpin loop adapter are designed to avoid regions of the construct where somatic hypermutations may occur. Framework (FR) and complementarity determining regions (CDR) regions are indicated within the figure. The CDR3 region arises from the juxtaposition of tandemly arranged variable, diversity and joining genes during V(D)J recombination.

For this demonstration (Methods), we designed two UMI-containing DNA probes targeting the terminal (invariable start and end) sequences, as well as a single UMI-containing stem-loop probe with feet complementary to the central linker domain of the chosen commercial phage display library (Figure 2). Once the UMI-containing full length library was diluted and amplified, mechanical fragmentation was carried out in such a way as to maximize the number of single cut fragments (Supplementary Figure 1). This is done to maintain high sample representation and to increase the probability of capturing UMI-containing reads; read-pairs without a UMI cannot be assigned to a putative original molecule and hence are discarded prior to assembly. Libraries were sequenced to saturation using the Singular Genomics G4™ platform and F2 flow cells (150M reads) with 2×150 cycle reads.

### Molecule reconstruction, filtering, and insert size distribution analysis

Post library sequencing, we first evaluated sample complexity by determining the number of reads associated with each detected UMI probe (Supplementary Figure 2, knee plot) following the approach in Melsted et al. 2019. This estimation is a good surrogate for the number of unique molecules in the library. As shown in Supplementary Figure 2, we detected 90,873 and 86,826 left probe and right probe associated UMIs, respectively, suggesting that the number of unique molecules was within this range. This number is roughly consistent with our expectation based on estimates from qPCR quantification, which indicated approximately 67,000 molecules were added to the PCR amplification reaction prior to fragmentation and adapter ligation. Given that the sample was sequenced to 47.8M paired-reads, we estimate the sequencing depth per input molecule to be within the range of 526 to 550 reads. At this sequencing depth we were able to assemble 58,170 contigs of >800bp in length, equivalent to 67% of the minimum estimated number of input molecules. This set of contigs might include members that only partially cover the expected VH and VL domains owing to errors occurring during the scFv library construction process. To remove such sequences from downstream analysis, we used IgBLAST (Ye et al. 2013) to extract the framework (FR) and complementary determining region (CDR) sequences for the VH and VL domains in each contig. Contigs having incomplete VH or VL FR1 or FR4 region sequences were filtered out, yielding 50,846 contigs containing full length VH-VL domains (mean length of 950bp; Figure 3 and Table 1).

**Table 1.** Reconstruction Metrics. Table indicates the estimated number of input molecules and the number of contigs satisfying various quality filters. Full length VH-VL contigs are defined as those containing complete VH and VL FR1 and FR4 region sequences, as determined by IgBlast annotation, and a minimum contig length of 800bp. Functional contigs are defined as those having VH and VL domains containing in frame variable and joining genes and no premature stop codons, as determined by IgBLAST.

| RECONSTRUCTION METRICS |  |
| --- | --- |
| Estimated Number of Input Molecules (UMI-based) | 86,826-90,873 |
| Number of contigs >800bp | 58,170 |
| Percentage input molecules reconstructed | 64-67% |
| Number of contigs containing full length VH-VL | 50,846 |
| Number of full length contigs having a stop codon | 1,411 |
| Number of full length contigs having an indel error | 3,937 |
| Number of full length, functional contigs | 45,498 |
| Percentage full length contigs with functional VH-VL | 89% |

**Figure 3.**
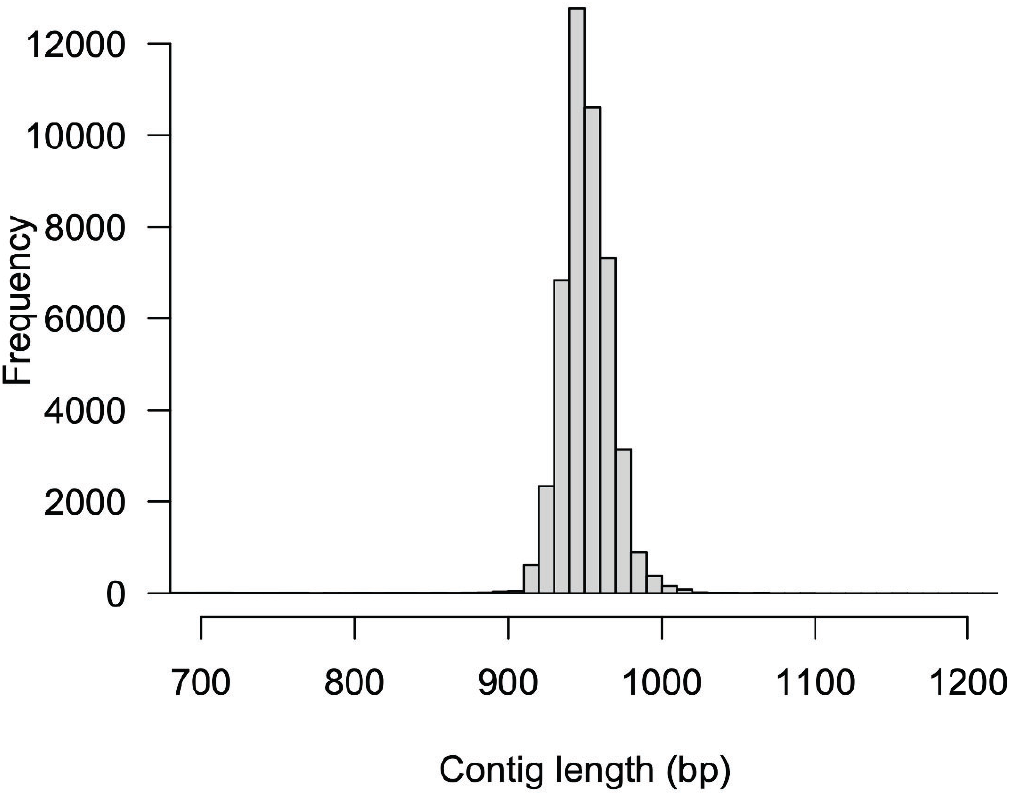
Size distribution of full length VH-VL contigs. Full length VH-VL contigs are defined as those containing complete VH and VL FR1 and FR4 region sequences, as determined by IgBlast annotation, and a minimum contig length of 800bp.

### VDJ analysis of VH-VL contigs

The VH and VL domains from assembled scFv contigs should reflect patterns of VDJ recombination and somatic hypermutation observed in studies using conventional high throughput sequencing to analyze heavy chain and light chain repertoires. To this end we examined variable gene and joining gene usage, CDR3 length distributions, and patterns of somatic hypermutation within the FR and CDR of reconstructed VH and VL rearrangements as determined by IgBLAST. We observe broad representation of variable and joining genes for both VH and VL rearrangements, with relative frequencies consistent with reports from literature (Jackson et al. 2013; Figure 4A). The frequency of IgK versus IgL containing contigs was 1:1.25, fitting expectation given the healthy donor material source. Heavy chain and light chain CDR3 length distributions match expectations for IgG expressing B cells (Figure 4 B), while somatic hypermutation analysis revealed elevated mutation of CDR vs FR regions (Murphy et al. 2008; Figure 4C). Further, we observe no correlation in the hypermutation level of paired VH-VL chains, as expected from random pairing of VH and VL chains during scFv library generation (Figure 4D). Finally, we observe high polyclonality, with a sample clonality of 0.001 (Methods), as expected from a diverse, unselected antibody display library.

**Figure 4.**
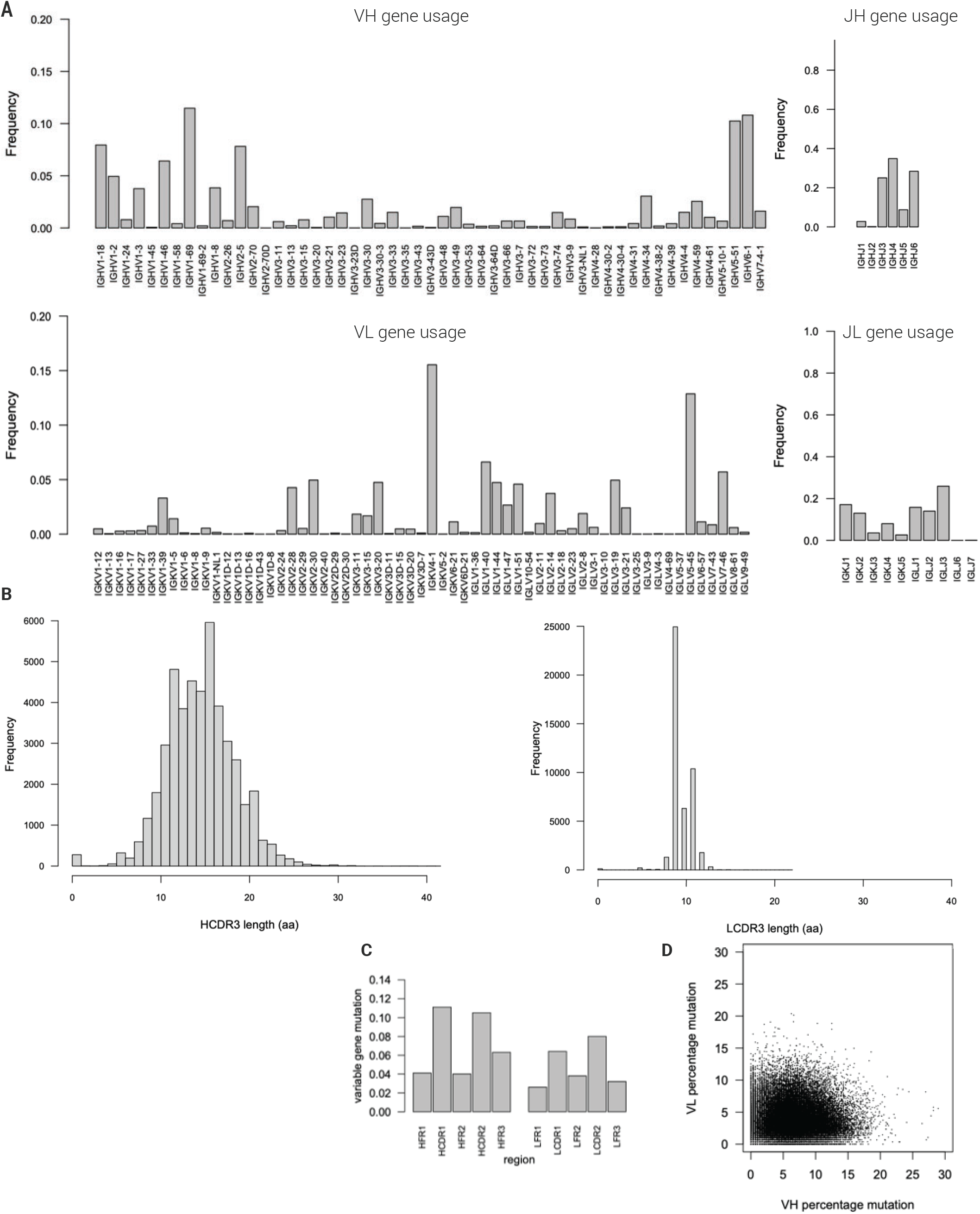
VDJ analysis of reconstructed VH-VL chains. A Variable gene and joining gene usage for VH and VL chains. Frequency indicates the fraction of VH or VL chains possessing a given variable or joining gene. B CDR3 length histograms for VH and VL chains. C Variable gene mutation rate as a function of IMGT annotated framework (FR) or complementarity determining regions (CDR). D Scatterplot of variable gene mutation rate for paired VH and VL chains. Each dot represents a unique VH-VL pair.

### Estimating the accuracy of reconstructed VH-VL contigs

One challenge in assessing the accuracy of reconstructed immune receptors is the lack of reference sequence with which to compare. To circumvent this issue, we estimated reconstruction accuracy by determining the fraction of full length VH-VL contigs having productive VH and VL rearrangements. Nonsense mediated decay and allelic exclusion should leave expressed immune receptors largely devoid of unproductive rearrangements (Murphy et al., 2008), an expectation that has been validated by next-generation sequencing of immune repertoires derived from RNA material (Warren et al. 2011). In our dataset, we find 89.5% (45,498) of full length VH-VL pairs to have productive VH and VL domains, with 2.8% of contigs having a premature stop codon in one of the chains and 7.7% having an insertion or deletion resulting in an out-of-frame rearrangement (Table 1). Taking into account the probability that a single base substitution or indel deriving from PCR or sequencing error gives rise to a premature stop codon or frameshift mutation, respectively, and the average length of the reconstructed VH-VL chains (Methods), we arrive at a minimum per base accuracy of 99.90% for reconstructed molecules, a figure that likely underestimates the true accuracy given that nonsense mediated decay and allelic exclusion do not completely eliminate unproductive rearrangements from RNA-based immune repertoire data (Lambert et al, 2020). Finally, as an independent method for determining accuracy, we calculated accuracy over the 45bp invariable linker region separating the VH and VL chains. Here, errors are readily identified by determining the Levenshtein edit distance between the observed and expected linker sequence for each full length VH-VL contig. Using this approach, we arrive at an accuracy of 99.91%, consistent with our estimate based on analysis of VH-VL productivity. In total, these results suggest an overall contig accuracy of approximately 99.9%.

### Assessing assembly efficiency as a function of sequencing depth

Lastly, we examined the relationship between sequencing depth and reconstruction efficiency, given that reconstruction efficiency is strongly affected by sequence accuracy and biases that may result in uneven coverage of template molecules (Alic et al. 2016, Heydari et al. 2017). Using downsampling analysis we find that ~100-200 reads per input molecule (~30-60X coverage) provides the optimal tradeoff between sequencing depth and contig yield (Figure 5, knee plot), as has been suggested elsewhere (Callahan et al. 2021).

**Figure 5.**
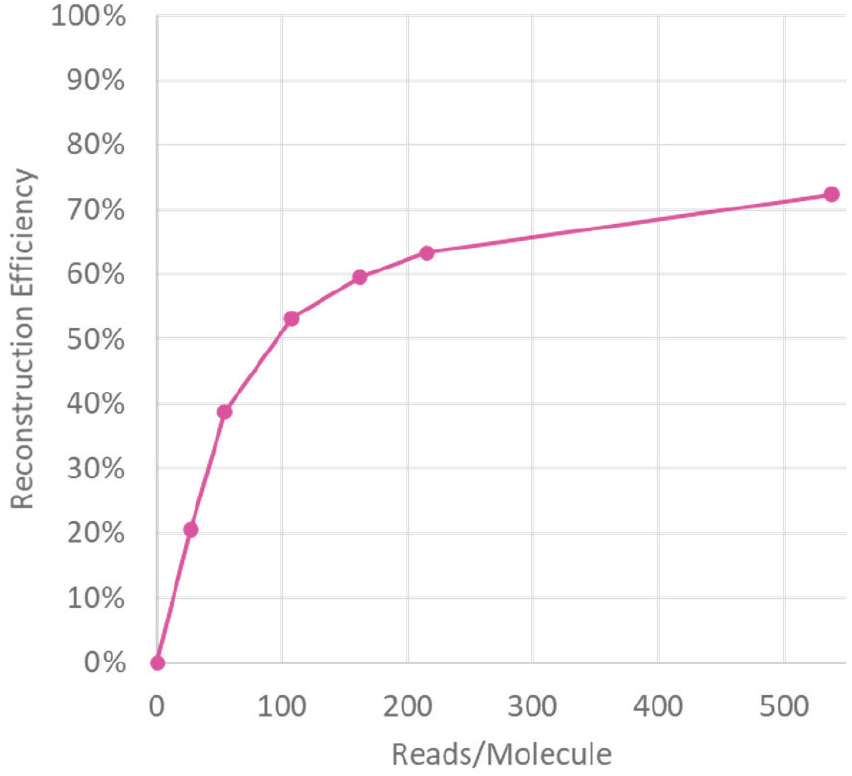
XR/T-Seq method assembly efficiency as a function of sequencing depth. The relationship between sequencing depth and assembly efficiency was assessed via downsampling of the dataset followed by contig assembly.

## Conclusions

The results presented here illustrate the utility of the Extended Range Targeted Sequencing method to characterize the properties of a commercial scFv Phage Display library. The XR/T-Seq method enables full length molecule reconstruction through the assembly of short reads associated into distinct UMI families. UMI tagging occurs through a primer extension and ligation reaction, which copies and uniquely labels the template DNA. The example presented here uses oligonucleotide probes that bind to specific locations within the template, where the distance between binding sites is critical for efficient read usage and UMI grouping. For 150bp paired reads and 1Kb templates, we designed probes to bind approximately 450bp apart and carried out fragmentation with conditions resulting in one cut or fewer per molecule. After paired-read 150 cycle sequencing and basic quality filtering, UMI tags are identified and grouped based off supporting read evidence. Read pairs are then grouped into the corresponding UMI families and assembled into final contigs using publicly available software.

The application of the XR/T-Seq method for characterizing scFv phage display libraries is ideal since these templates have a perfectly suited size range, 0.8-1Kb, and have fully conserved regions introduced during the manufacturing process. With this model system we were able to reconstruct approximately 65% of the input molecules: 58,170 contigs larger than 850bp, out of an estimated 86,826-90,873 input molecules. This maximum number of reconstructed templates was achieved at 538 reads per molecule (using all reads), though downsampling analysis revealed approximately 30-fold coverage as the optimal sequencing depth per molecule, with an extrapolated throughput of ~600K full length reconstructed scFv fragments per F2 flow cell (150M reads capacity). Analysis of VH-VL pairs demonstrates that library composition fits expectation from random pairing of heavy and light chains derived from healthy donor IgG expressing B cells, with variable, diversity and joining gene representation and somatic hypermutation profiles matching expectations from other immune repertoire sequencing studies. Finally, the use of short reads for assembly results in high overall accuracies, here estimated to be approximately 99.9%.

We expect this method to be useful for other applications where targets of interest are in the 1-3Kb range, and where appropriately spaced probes can be designed to target conserved regions (e.g., 16S RNA profiling, quantifying tandem repeat expansions, or characterizing human leukocyte antigen (HLA) regions). For example, 16S sequencing has become popular among microbiologists as a method to understand the diversity and population structure of microbes in specific ecosystems. Due to limitations in short read NGS platforms, the entirety of the 16S gene (approximately 1,500-1,800 bp) is otherwise difficult to accurately sequence. In summary, we expect the XR/T-Seq method to aid in the characterization of complex DNA libraries with high accuracy and throughput.

## Methods

### ScFv Template Generation

The template used in this study was derived from a commercial single chain phage display naïve human antibody library consisting of randomly paired Variable Heavy (VH) and Variable Light (VL) domains of IgG expressing B cells independently amplified from a pool of healthy donors. Paired VH-VL domains were created in an expression vector by inserting the following configuration: *Invariable Start-VH-Linker-VL-Invariable End*, where *Linker* comprises a 45bp invariant sequence encoding a flexible linker region composed of glycine and serine residues.

*Invariable Start* and biotinylated *Invariable End* primers (0.2uM each) were used to amplify an aliquot of the parent library (20ng per 50μL reaction) with 16 cycles of PCR (1× Phusion HF Buffer, 1U Phusion U Polymerase, dNTP mix (2.5nM dATP, 2.5nM dGTP, 2.5nM dCTP, 1.25nM dTTP, 1.25nM dUTP)). Phusion U Polymerase (Thermo) was used to randomly incorporate dUTPs into the template for a downstream digestion of the template. PCR amplicons were purified with QuantaBio’s sparQ PureMag beads (sparQ beads) using a 0.8x ratio and quantified by fluorescence (Qubit)

### Extension and Ligation

Biotinylated template was immobilized by binding to streptavidin beads. Briefly, 10 μL of My1C1 beads (Invitrogen) were added to a 1.5mL tube, placed on a magnetic rack to remove buffer, washed twice with 1× binding buffer (aqueous solution including Tris-HCl, NaCl, EDTA and Tween-20), and resuspended in 2× binding buffer to a final volume of 20μL. 0.25pmol (152ng) of biotinylated template adjusted to 20μL using nuclease-free water was added to the bead suspension and incubated at RT for 1 hour with frequent mixing (rotation). At the completion of this binding step, the tube was placed on a magnetic rack, supernatant removed, and beads washed twice with 1× binding buffer (200μL/wash). Beads were then resuspended with 10μL 0.1M NaOH and incubated at RT for 3 minutes. The tube was then transferred to a magnetic rack and supernatant discarded. Beads were washed twice by resuspending in 200μL 1× binding buffer. After discarding supernatant at the end of the second wash, beads were resuspended in 14μL bead master mix (1× T4 DNA Ligase Buffer (NEB), 125uM each dNTP, 1× TE pH 8.0). Pooled (x3) UMI-containing probes at 12.5uM each (1μL each) were 5’ phosphorylated by T4 Polynucleotide Kinase (NEB) (1× T4 DNA Ligase Buffer, 2.4U PNK) directly prior to hybridization. Adapters were added to immobilized template for a final volume of 46μL per reaction. Tubes were transferred to a thermal cycler for hybridization (10sec at 90°C, ramp down to 37°C at −0.2°C/s, and incubation at 37°C for 30min) and immediately supplemented with 1μL T4 DNA Pol (40,000 U/μL, NEB), 3μL of T4 DNA Ligase (20,000 U/μL, NEB) to 50μL final volume. The tube was further incubated at 37°C for 1 hour. Reactions were cleaned up by placing the strip tube on a magnetic rack, discarding supernatant, and washing beads twice with 200μL 1× binding buffer. Extended-ligated products were eluted from beads with a 3min incubation at RT in 20μL 0.1M NaOH. Supernatant was transferred to a fresh tube with 9μL 200mM Tris-HCl, pH 7.0 to neutralize the sodium hydroxide. Extended-ligated products then underwent USER digestion (1× Cutsmart Buffer, 1U Thermolabile USER II (NEB)) to remove template, purified with sparQ beads using a 2× ratio, visualized on a 6% TBE-Urea denaturing gel (120 min at 220V), and quantified by qPCR.

### Tagged-DNA Amplification

Input molecules were amplified by 25 cycles of PCR using external PCR primers, designed to amplify full length molecules. Full-length DNA molecules were purified with sparQ beads and quantified by fluorescence (Qubit).

### NGS Library Preparation

112ng of full-length UMI-tagged DNA molecules were fragmented using a Covaris ME220 Sonicator, with 3 iterations of the following treatment: Duration = 40.0 s; Peak Power = 50.0; Duty% Factor = 10.0; Cycles/Burst = 1000; Avg Power = 5.0. PCR-Free libraries were made with an adapted version of QuantaBio’s sparQ DNA Library Prep kit, where ligation was performed with adapters containing full length sequencing primer binding regions and Singular Genomics’ clustering sequences. Libraries were size selected to mean insert sizes ranging from 450 to 600bp using sparQ beads for size selection.

### Sequencing

Libraries prepared according the XR/T-Seq method were sequenced on F2 flow cells using Singular Genomics’ G4 platform. Briefly, libraries were diluted to 25pM in loading buffer and loaded into the sample cartridge. The loaded libraries were amplified and sequenced using a paired-read 150 cycle format at sufficient depth to allow for >500 paired-reads per original molecule.

### Reconstruction of UMI-tagged Molecules

The assembly process consists of three steps. First, UMIs are extracted from sequence reads and sequencing or PCR errors removed by collapsing sequences within an edit distance of 1 via UMIcollapse (https://github.com/Daniel-Liu-c0deb0t/UMICollapse). Next, network analysis (via Python NetworkX library) is used to identify co-occurring groups of UMIs derived from the same starting molecule, with low confidence connections (edges) removed on the basis of raw read counts and the proportion of reads for a given UMI supporting a given edge. Finally, UMI groups are used to partition reads into read clusters, each of which is subjected to K-mer based error removal followed by SPAdes assembly (Bankevich et al. 2012).

### VDJ Analysis of VH-VL Contigs

Following assembly, VH and VL domains of assembled contigs were analyzed via IgBLAST (Ye et al. 2013) to annotate rearrangement features. Full length contigs (defined as contigs having complete FR1 and FR4 regions of both VH and VL rearrangements and a contig length >=800bp) were identified and used to assess variable, diversity, and joining gene usage, CDR3 length distributions, somatic hypermutation, and rearrangement productivity.

### Calculation of Clonality

Clonality was calculated by first determining the frequency of each clonotype in the dataset, where a clonotype is defined as a contig having a unique combination of VH gene identity, VL gene identity, HCDR3 and LCDR3 sequence. Clonality was then defined using the Python (v3.8.6; scipy v1.5.2) formula:

Clonality = 1-scipy.stats.entropy(frequencies, base=2)/math.log(len(frequencies),2)

where *frequencies* is the list of clonotype frequencies in the dataset.

### Estimating Contig Accuracy

To estimate the sequence accuracy of reconstructed VH-VL chains, we determined the frequency of full length contigs having either a frameshift mutation or premature stop codon resulting in an unproductive VH or VL domain. We then used the following assumptions to estimate the per base error rate:

1. Allelic exclusion and nonsense mediated decay are totally efficient such that all VH and VL chains in the RNA-derived scFv library are productive.
2. All single base insertion or deletion errors should give rise to a frameshift mutation.
3. Only a subset of single base substitution errors will convert an arbitrary codon into a premature stop codon given that only a subset of codons (18 of 61) are within a Hamming distance of 1 from a stop codon. More precisely, only 23 of the 61*9 potential single base substitution events will convert a non-stop codon into a stop codon. Thus, the observed frequency of premature stop codon containing rearrangements likely underestimates the true number of substitution errors in the data. To account for this, and further assuming that all codons are found at approximately equal abundance in the data, we multiply the number of detected premature stop codons by (61*9/23).

Applying these assumptions, and given an average combined VH-VL length of 739bp (excluding the invariant linker region and adapter sequences), we define the per base accuracy as:

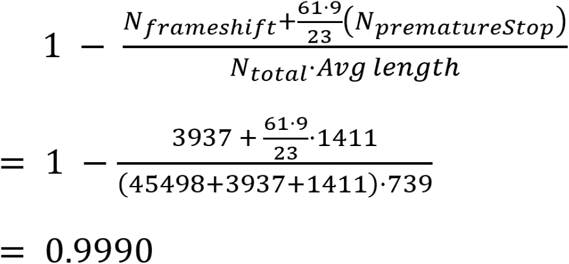

Accuracy over the 45bp linker region was calculated by determining the sum of the Levenshtein edit distances between the observed and expected linker sequence for each full length VH-VL contig, then dividing this number by the total number of linker bases analyzed.

