## Supplementary Figures for "Reconstruction of Full-length scFv Libraries with the Extended Range Targeted Sequencing Method"

**Supplemental Figures**

Christopher Wei, Sarah Criner, Bharat Sridhar, Ryan Shultzaberger, Timothy Looney, Martin M Fabani, Eli N Glezer

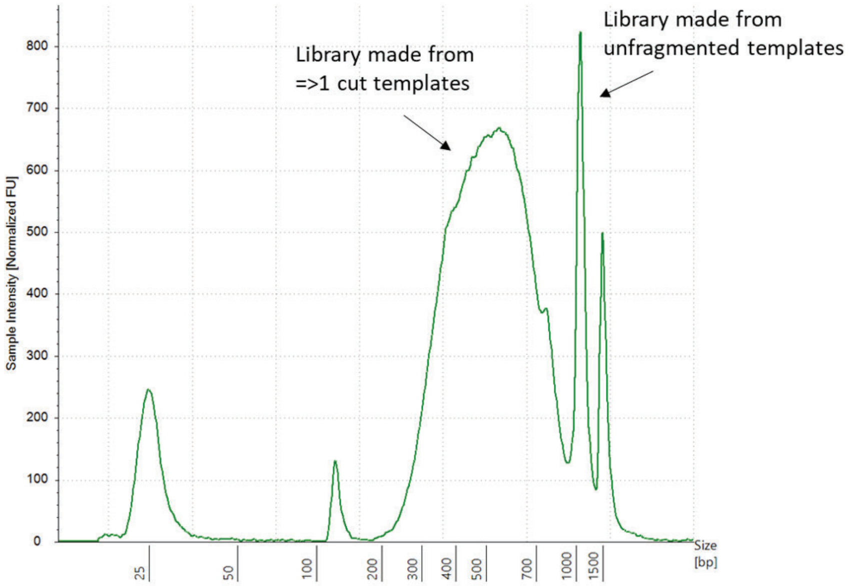


**Supplementary Figure 1**: Library size distribution. Size distribution profile for representative library obtained using fragmentation and size selection conditions targeting 450-600bp in size (including sequencing adaptors).

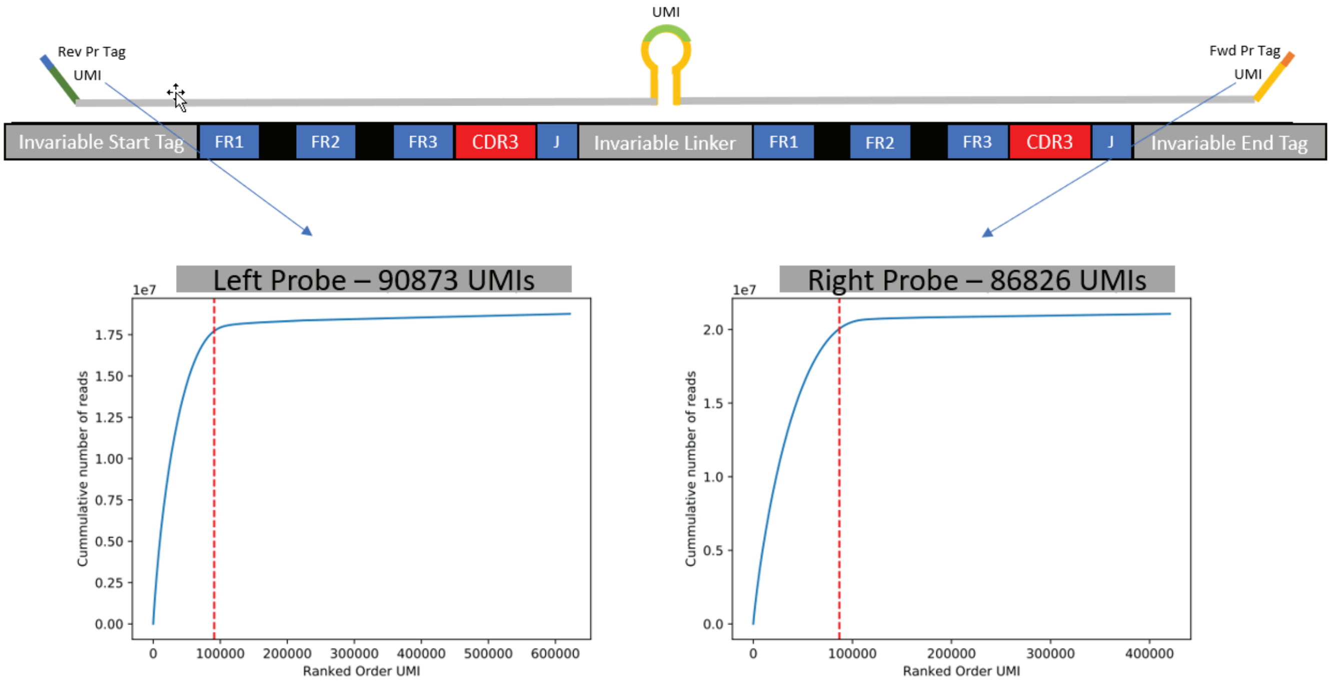


**Supplementary Figure 2**: Estimation of input molecules by UMI counting. Input molecule number was estimated by determining the cumulative number of reads for unique UMIs.
